# Volumetric mechanosensing of CAF in 3D hydrogels drive altered drug response in breast cancer

**DOI:** 10.64898/2026.02.25.707983

**Authors:** Somayadineshraj Devarsou, Nam Ji Sung, Seok Hoon Ham, Martin Kiwanuka, Jennifer L. Young, Jennifer H. Shin

## Abstract

Altered mechanical properties of the tumor microenvironment (TME) influence cancer progression, yet the mechanistic basis by which 3D mechanics shape CAF heterogeneity and downstream tumor drug response remains poorly understood. Here, we engineered a modulus-tunable gelatin methacryloyl (GelMA) hydrogel platform spanning a normal-like (soft ∼2 kPa) to desmoplastic-like (stiff∼40 kPa) range to culture primary breast CAFs under 3D confinement. CAFs exhibited pronounced volumetric morphoadaptation across matrices, with soft 3D matrices supporting larger, more protrusive morphologies and stiff gels constraining cell geometry. In contrast to canonical 2D paradigms, nuclear YAP localization was reduced in stiff 3D matrices and varied substantially across cells, consistent with a dominant role for 3D geometric/volumetric state in regulating mechanotransduction. Functionally, in transwell co-culture with MCF-7 spheroids under paclitaxel treatment, CAFs cultured in stiff 3D matrices induced a broader chemoresistance-associated transcriptional program, whereas soft 3D matrices CAFs favored stress/checkpoint-like responses. A 2D monolayer comparator indicated that coordinated resistance-associated programs emerge most clearly in 3D tumor architecture. Together, these results establish a GelMA-based biomaterials framework in which CAF volumetric state provides a quantifiable intermediate linking 3D matrix mechanics to mechanotransduction and tumor drug-response programs, motivating future strategies to modulate stromal function through mechanically controlled cell-state regulation.

## 1. Introduction

Cancer-associated fibroblasts (CAFs) are increasingly recognized as pivotal drivers of the tumor microenvironment (TME), mediating biochemical and biophysical cues that critically shape malignant cell fate and therapeutic response [1–6]. The functional diversity of CAFs stems from pronounced phenotypic heterogeneity, dynamically regulated in space and time through interactions with tumor cells, the extracellular matrix (ECM), and other stromal components [1,7–9]. This heterogeneity poses a major obstacle to the development of effective CAF-targeted therapies. Among the defining features of the TME, biomechanical cues derived from the ECM play a central role in governing cell morphology, mechanotransduction, differentiation, and intercellular communication [5]. ECM stiffness has emerged as a potent regulator of both cancer cell behavior and stromal cell function [10,11].

CAFs, by virtue of their mesenchymal origin, are highly sensitive to mechanical cues within the TME [12–14]. Breast cancer, the most prevalent malignancy among women and a disease frequently associated with therapeutic resistance, is characterized by progressive ECM stiffening driven by desmoplasia, a fibrotic remodeling process marked by excessive collagen deposition [15,16]. Prior studies have shown that matrix stiffness can modulate CAF phenotype in 3D hydrogels [17–20], but how stiffness-dependent CAF volumetric adaptation links mechanical cues to paracrine signaling and tumor drug responses remains unclear.

Despite this recognition, it remains unclear whether a single, quantifiable biophysical parameter, such as ECM stiffness, can systematically stratify CAFs into discrete, functionally targetable states that account for their pronounced heterogeneity in tumor progression and chemoresistance [21–24]. To test this hypothesis, we engineered a three-dimensional (3D), stiffness-tunable GelMA hydrogel platform spanning physiologically relevant low-modulus (soft, normal-range, ∼2 kPa) and high-modulus (stiff, desmoplastic-range, ∼40 kPa) states to recapitulate the mechanical transition from normal to desmoplastic tumor ECM.

We show that CAFs undergo stiffness-dependent morphoadaptation: in soft GelMA matrices, cells expand in volume and exhibit more protrusive, irregular 3D morphologies, whereas in stiff matrices, cells become volumetrically restricted and adopt more compact configurations. These volumetric states are accompanied by distinct transcriptional profiles. For example, CAFs in stiff matrices show the upregulation of PDPN, ACTA2, and MMP2, as well as preferential engagement of MRTF-A and TRPV4 signaling, whereas in soft matrices, CAFs maintain a less contractile state characterized by elevated nuclear YAP/TAZ activity. While ECM stiffness is widely viewed as a dominant regulator of fibroblast activation through YAP-dependent mechanotransduction [25], this framework is largely derived from 2D systems and does not translate to 3D environments [26–30]. It remains unclear how activated CAFs persist in stiff matrices, where canonical YAP nuclear localization is suppressed. Here, we focus on the CAF volumetric state (cell volume and shape) under 3D bulk modulus as a quantifiable intermediate linking ECM mechanics to downstream transcriptional responses.

Functionally, CAFs cultured in stiff (vs. soft) GelMA matrices differentially regulate tumor drug-response programs. Specifically, CAFs cultured in stiff matrices promote a chemoresistance-associated transcriptional program in MCF-7 spheroids, marked by increased gene expression of ABCB1, BIRC5, EGFR, and MMP2, whereas in soft matrices, CAFs favor stress-adaptive programs associated with CDKN2A (p16) and CDKN1A (p21). Consistent with this functional divergence, cytokine profiling revealed that CAFs in stiff matrices, secrete higher levels of IL-6, MCP-1, OPG, and TIMP-1, reinforcing a chemoprotective tumor microenvironment. Together, these results suggest that the stiffness-dependent volumetric state of CAFs, defined by cell size and shape under 3D confinement, serves as a measurable intermediary between ECM mechanics and tumor drug-response programs.

## 2. Methods

### 2.1. Cell culture

Primary breast cancer–associated fibroblasts (CAFs) isolated from invasive breast adenocarcinoma tissue (Bio IVT) were used for the entire analysis. CAFs were cultured in Dulbecco’s Modified Eagle Medium (DMEM) supplemented with 10% fetal bovine serum (FBS) and 1% penicillin–streptomycin (PS). Cultures were maintained at 37 °C in a humidified incubator with 5% CO₂, with medium changes every 2 days. MCF-7 human breast cancer cells (ATCC) were maintained in DMEM supplemented with 10% FBS and 1% PS under standard culture conditions (37 °C, 5% CO₂).

### 2.2. GelMA Synthesis

Gelatin methacryloyl (GelMA) was synthesized by methacrylation of gelatin following a previously reported protocol[28]. Briefly, 10 g gelatin powder (Porcine gelatin, Type A, 300 bloom) was dissolved in 100 mL phosphate-buffered saline (PBS) at 60 °C for 1 h. Methacrylic anhydride (Sigma Aldrich) – 8 mL was added dropwise under vigorous stirring, and the reaction proceeded for 2 h at 60 °C. An additional 100 mL of prewarmed PBS was added to terminate the methacrylation process, and the solution was maintained at 60 °C for 30 min. The product was dialyzed against deionized water at 40 °C for 7 days, lyophilized, and stored at −20 °C.

### 2.3. GelMA Hydrogel Fabrication

GelMA hydrogel matrices (10% w/v) were prepared by dissolving 0.1 g GelMA in 0.980 mL PBS at 40 °C for 30 min. Cell suspensions were added to achieve a density of 150,000 cells per gel and gently homogenized. Crosslinking was initiated using a visible-light photoinitiator system containing 0.5 mM ruthenium and 5 mM sodium persulfate. GelMA was cast in 6-well plates (for RNA analysis) or confocal dishes (for imaging) and polymerized under visible light. Exposure times were optimized to generate low-modulus (soft, normal-range, ∼2 kPa) and high-modulus (stiff, desmoplastic-range, ∼40 kPa)

### 2.4. CryoSEM

To preserve native pore architecture, GelMA was cryo-fixed in liquid nitrogen and imaged using a multifunctional cryo-focused ion beam system (Aquilos/ Thermo Fisher Scientific/ KARA - KAIST Analysis Center for Research Advancement, KAIST). Samples were maintained at cryogenic temperatures throughout imaging to prevent structural deformation.

### 2.5. Rheological Characterization

Viscoelastic properties of GelMA were measured using a rheometer with parallel plate geometry. Oscillatory stress sweep tests were performed under small-amplitude shear to determine storage modulus (G′) and elastic behavior.

### 2.6. Confocal Imaging

Confocal laser scanning microscopy was performed using a Carl Zeiss LSM 980 KARA (KAIST Analysis Center for Research Advancement, KAIST). 3D Z-stack imaging was conducted to capture 3D cellular morphology, with imaging depth and step size adjusted according to sample thickness. Z-stacks were collected at a fixed depth (30-50μm) from the gel surface across all samples.

### 2.7. 3D image analysis

Volumetric segmentation and morphometric analysis were performed using Imaris software. Extracted features included total cell volume, surface area, sphericity, and elongation ratio. Protein localization was quantified within reconstructed 3D cell models. For each segmented cell, static 3D morphometric features were extracted using Imaris. These included total cell volume, surface area, sphericity, aspect ratio, circularity, convex hull metrics, and a contractility index derived from shape irregularity. These parameters quantified the instantaneous volumetric and geometric state of each CAF within the 3D matrix.

YAP activity was quantified using a nuclear-to-cytoplasmic (N/C) ratio calculated as

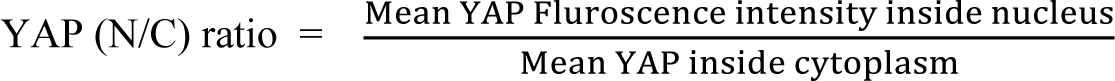

Nuclear and cytoplasmic regions were defined using the segmented nuclear mask and the remaining cytoplasmic cell volume, respectively. Mean fluorescence intensities were computed within these compartments for each cell.

PDPN expression was quantified as the total integrated fluorescence intensity within the segmented 3D cell volume. PDPN intensity distributions were correlated with cell volume and other morphometric features to assess relationships between volumetric state and mechanosensitive marker expression.

### 2.8. Machine learning based classification of CAF mechanical states

The standardized feature matrix was used as input for supervised machine learning classifiers and trained to distinguish CAF mechanical states in soft versus stiff GelMA matrices.

Model outputs were visualized using dimensionality reduction techniques (UMAP and PCA), confusion matrix, and box plots to assess classification performance and feature separability. Feature importance scores were used to compare the predictive power of static morphometric features versus dynamic biophysical features.

### 2.9. MCF-7 spheroid generation and co-culture

MCF-7 cells were seeded at 5,000 cells per well in ultra-low adhesion plates. Collagen I (6.25 mg/mL) was added to promote spheroid compaction. Spheroids were cultured for 4 days prior to co-culture. CAFs were encapsulated in soft ∼2 kPa and stiff ∼40 kPa, GelMA matrices and cultured for 4 days. CAF-laden GelMA were placed in the upper compartment of a transwell-style system, with MCF-7 spheroids maintained below, enabling paracrine-only signaling.

### 2.10. Drug treatment and viability assay

Co-culture experiments were treated with paclitaxel (6.25 ng/mL) for the indicated duration, with DMSO (dimethyl sulfoxide) used as the vehicle control across all conditions. Spheroid viability after drug treatment was assessed using a Calcein-AM/propidium iodide (PI) Live/Dead assay (MaxView BDA-1000).

A fresh working stain solution was prepared on the day of use by mixing 5 mL D-PBS with 10 μL Solution A (Calcein-AM stock) and 15 μL Solution B (PI stock), protected from light. For staining, culture medium was gently removed without aspirating spheroids, spheroids were washed once with D-PBS, and then incubated in FBS-free medium containing the working stain solution at 37 °C for 30 min. Spheroids were optionally washed once with D-PBS to reduce background and then imaged by widefield/epifluorescence microscopy, focusing on the mid-plane and edge regions. Calcein-AM (live) and PI (dead) fluorescence were recorded under identical settings across conditions.

### 2.11. RNA isolation and qRT-PCR

GelMA matrices were digested using collagenase type II (2 h). RNA was isolated and analyzed using the Bioneer AccuTarget™ qPCR Screening Kit (20-SH-0001-10-CFG), which includes pre-validated primer sets for key human breast cancer targets. Complementary DNA (cDNA) Synthesis and quantitative real-time PCR were carried out according to the manufacturer’s instructions. Relative gene expression levels were calculated using the ΔΔCt method, allowing for comparative analysis between experimental groups. Relative expression was calculated using the ΔΔCt method, normalized to GAPDH.

### 2.12. Cytokine Array Analysis

Cell-free conditioned media (CM) were collected from CAFs in soft and stiff GelMA matrices at day 7 to analyze the secreted protein profiles. Cytokine expression was evaluated using the Human Cytokine Antibody Array (membrane-based, 80 targets; Abcam, ab133998). Chemiluminescence signals were captured and quantified, and protein levels were normalized against the array’s internal positive controls. This analysis enabled the identification of differential cytokine secretion patterns associated with matrix stiffness-dependent CAF states.

### 2.13. Statistical analysis

Data presented in figures are expressed as mean ± SEM unless stated otherwise. Statistical comparisons were performed using Student’s t-test or one-way ANOVA with post-hoc testing. Significance thresholds were *p < 0.05, **p < 0.01, ***p < 0.001. All experiments were performed in triplicate or more unless otherwise noted, and in such cases, the number of cells used in the measurement has been stated. Analyses were performed using Origin and Microsoft Excel.

## 3. Results

We used a 3D GelMA hydrogel platform with tunable mechanical properties to model the transition from a soft to a stiff ECM, matching healthy (1.3–2.4 kPa) to desmoplastic invasive ductal adenocarcinoma (30-40 kPa) [31]. GelMA was synthesized and characterized by ¹H NMR and FTIR (Fig. S1), confirming methacrylation suitable for Ru/SPS visible-light crosslinking (DoM = 63%). To tune the elastic modulus, we varied the photo-crosslinking exposure time (0.5 min for low-modulus- soft range ∼2 kPa; 9 min for high-modulus - stiff range ∼40 kPa) with minimal cytotoxicity (<20%) (Fig. S8). These formulations produced reproducible differences in both mechanics and microstructure: soft matrices exhibited a more open network with larger pores, whereas stiff matrices formed a denser network with smaller pores (Fig. S2), consistent with the measured elastic modulus (Fig. 1B). Because modulus was tuned by photocrosslinking, increases in bulk stiffness were accompanied by changes in network density and microarchitecture. Thus, the observed volumetric restriction likely reflects the combined effects of increased confinement and altered network structure, rather than modulus alone.

**Fig. 1.**
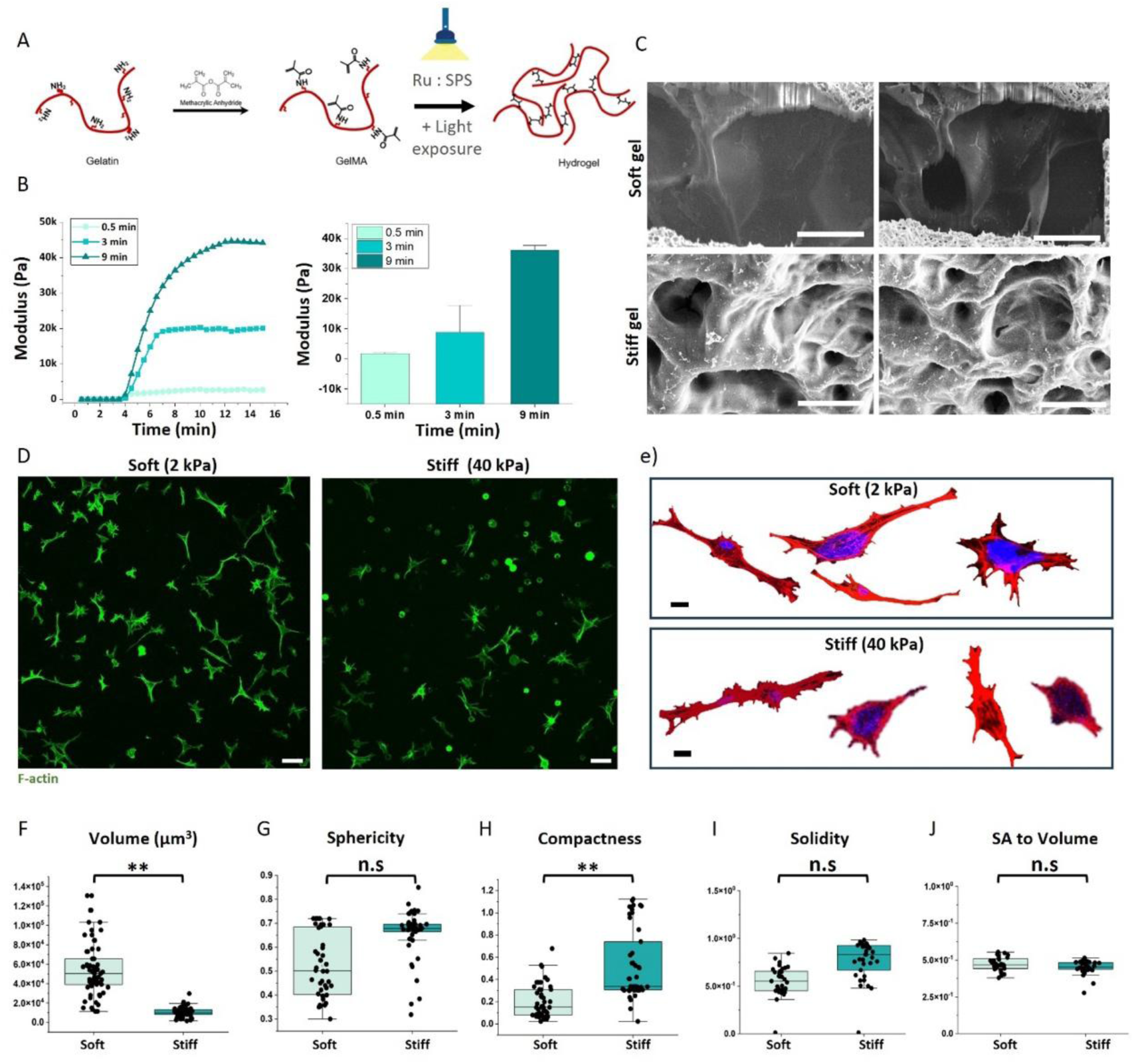
Engineering a tunable 3D GelMA hydrogel platform to control matrix compliance and CAF volumetric adaptation. A) Schematic of GelMA synthesis via methacrylation of gelatin and subsequent photo-crosslinking using the Ru/SPS initiator system. Light exposure induces covalent network formation, generating 3D GelMA hydrogels with tunable mechanical properties. B) Rheological characterization showing time-dependent increases in storage modulus (G′) during light exposure. Increasing crosslinking duration yields progressively stiffer matrices, enabling tuning from ∼2 to ∼40 kPa. C) Representative scanning electron microscopy (SEM) images of GelMA hydrogels crosslinked to soft (∼2 kPa) and stiff (∼40 kPa) matrix conditions. Soft gels exhibit larger, more open pores, whereas stiff gels display a denser, more compact microarchitecture. Scale bar, 5 μm. D) Representative confocal images of CAFs encapsulated in soft (2 kPa) and stiff (40 kPa) GelMA matrices and stained for F-actin (green) and nuclei (DAPI, blue). Scale bar, 100 μm. E) Representative single-cell 3D segmentation of CAF morphologies in soft and stiff GelMA matrices (F-actin, red; nuclei, blue). Scale bar, 10 μm. F-J) Quantitative 3D morphometric analysis from single-cell reconstructions showing matrix-dependent differences in total cell volume, sphericity, compactness, solidity, and surface area (SA)-to-volume ratio. Each point represents one cell (n ≈ 30–35 cells per condition; n ≈ 3–5 gel conditions). Box plots indicate median and interquartile range. Statistical significance was assessed by Student’s t-test (*p < 0.05, **p < 0.01).

Primary breast CAFs were encapsulated in soft ∼2 kPa and stiff ∼40 kPa GelMA matrices and cultured for 4 days. Morphological assessment from 3D reconstructions revealed stiffness-dependent differences, with CAFs in soft matrices exhibiting larger cell volumes, greater surface area, and more rounded morphologies than those in stiff matrices (Fig. 1D, E). To quantify these changes, we used a minimal set of robust global 3D morphometrics (volume, surface area, sphericity, elongation, and compactness) that operationalize CAF “volumetric state” and are comparatively less sensitive to surface-mesh artifacts than curvature- or skeleton-derived descriptors. As a complementary measure of protrusive, non-convex geometry, we also computed solidity (volume/convex-hull volume), which provides an interpretable readout of branching/spikiness while remaining stable under a moderate smooth surface. GelMA concentration (10%) and the biochemical composition of the culture media (10% FBS, DMEM) were held constant across conditions (1.3 × 10^5^ cells/gel). Accordingly, the observed differences in CAF morphology are attributable primarily to the hydrogel mechanical state, soft versus stiff GelMA matrices. These results suggest that the stiff gels constrain CAF volumetric expansion consistent with increased mechanical resistance to isotropic growth, whereas soft gels permit greater volumetric expansion. However, because morphology was quantified at Day 4, substantial cell-to-cell heterogeneity within each stiffness condition suggests that local microstructural variability and/or time-dependent matrix remodeling may also contribute.

To assess whether stiffness-dependent volumetric adaptation of CAFs is accompanied by coordinated changes in functional phenotype, we quantified CAF gene expression by reverse transcription quantitative PCR (RT-qPCR). To contextualize stiffness-dependent gene expression changes observed in 3D, we included 2D gelatin-coated substrates as a reference condition representing conventional CAF culture without 3D stiffness. We then compared gene expressions across 2D and 3D soft (vs. stiff) conditions (Fig. 2A). Consistent with these findings, immunostaining revealed elevated PDPN protein expression in CAFs cultured in stiff matrices, accompanied by a polarized actin cytoskeleton (Fig. 2B). To clarify the basis for CAF phenotypic differences, we grouped stiffness-regulated genes into functional categories. Notably, CAFs in stiff matrices showed selective upregulation of matrix secretion and remodeling signatures, rather than a uniform increase across canonical myofibroblastic markers (Fig. 2C), indicating persistent phenotypic heterogeneity even within stiffness-defined CAF populations.

**Fig. 2.**
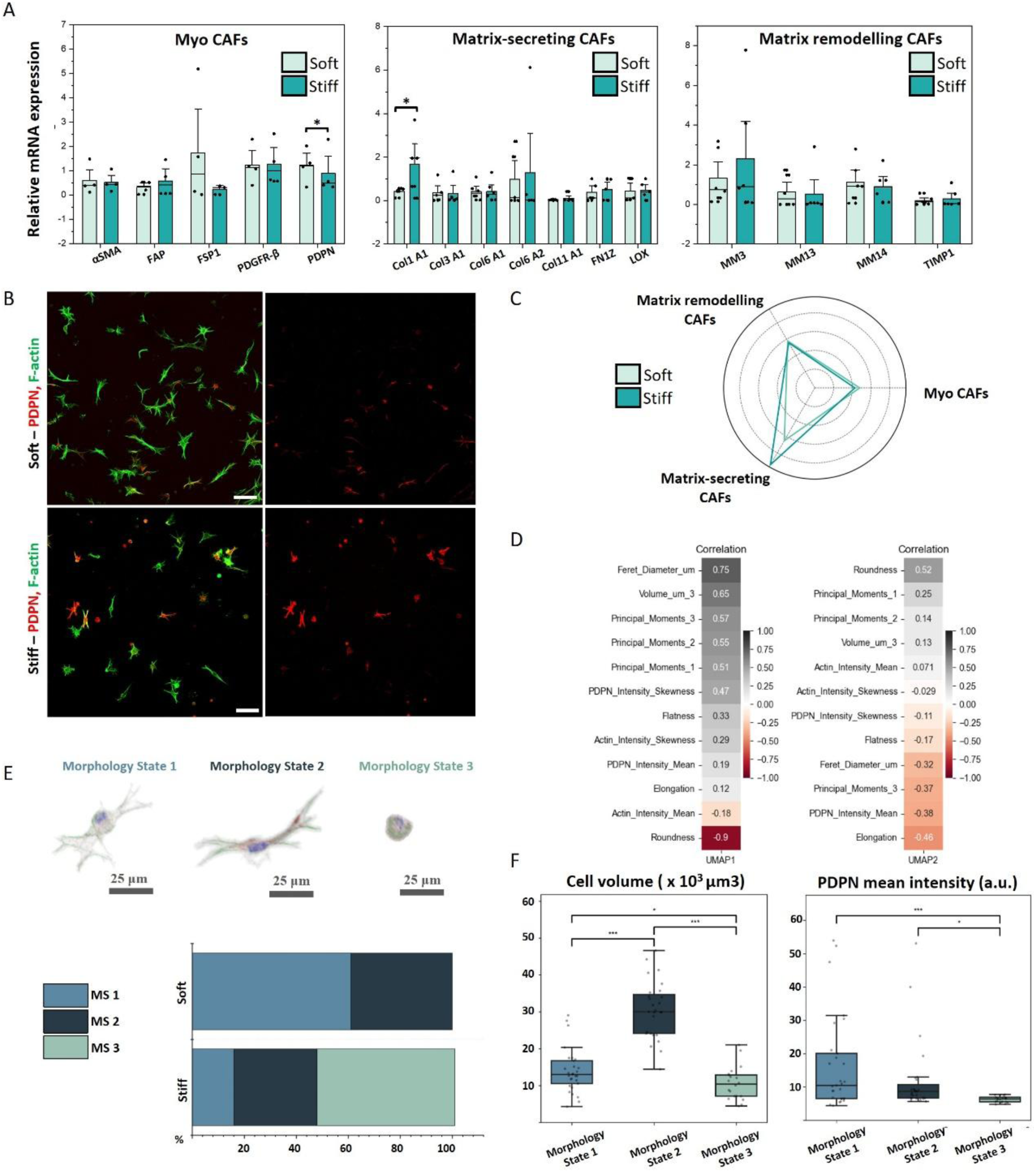
Soft versus stiff 3D matrices program CAF transcriptional identity through discrete volumetric and morphological states. A) Quantitative RT–qPCR analysis of CAF subtype-associated gene expression in soft (∼2 kPa) and stiff (∼40 kPa) GelMA matrices. Genes associated with myofibroblastic CAFs (myCAFs), matrix-secreting CAFs, and matrix-remodelling CAFs are shown. Expression values are normalized to 2D CAF controls and presented as relative mRNA expression. Bars represent mean ± SD; individual dots indicate biological replicates (n = 4–6). Statistical significance was assessed by one-way ANOVA with post hoc testing (*p < 0.05, **p < 0.01). B) Representative immunofluorescence images of CAFs cultured in soft (2 kPa) and stiff (40 kPa) GelMA matrices, stained for F-actin (green) and podoplanin (PDPN, red). Scale bar, 100 μm. C) Radar plot summarizing normalized functional enrichment scores for matrix secretion, matrix remodelling, and myo CAF-associated programs in CAFs cultured in soft versus stiff matrices. D) Heatmap showing correlations between principal morphometric features and UMAP dimensions derived from single-cell volumetric analysis. E) Representative 3D reconstructions illustrating three distinct CAF morphological states identified by unsupervised clustering. Scale bar, 25 μm. The heatmap below shows the distribution (%) of each morphology state across soft and stiff matrix conditions. F) Quantification of cell volume and elongation across morphology states. Box plots indicate median and interquartile range; whiskers denote minimum and maximum values. Statistical significance was determined by one-way ANOVA (*p < 0.05, **p < 0.01). PDPN, podoplanin; UMAP, uniform manifold approximation and projection.

To determine whether these stiffness-dependent morphological features resolve discrete single-cell states, we used a deep learning-based pipeline to reconstruct >300 CAFs in 3D and extract morphometric parameters for clustering analysis. Three distinct morphological states emerged, primarily differentiated by elongation, roundness, and total cell volume (Fig. 2D, E). Compact, low-volume forms dominated in stiff matrices, whereas elongated forms were common in soft matrices, and no morphological state was exclusive to any stiffness.

Next, we asked whether CAF morphological states are associated with marker expression at the single-cell level. Correlation analysis indicated that PDPN expression varied most strongly with specific morphometric features, particularly elongation parameters, rather than with the bulk modulus of the GelMA alone (Fig. 2F). Density-based analysis of segmented 3D reconstructions further visualized continuous morphological transitions within the CAF population (Fig. S3).

Collectively, these findings suggest that CAF volumetric state, defined by changes in cell volume and shape under 3D confinement, serves as a quantifiable intermediary linking ECM stiffness to downstream transcriptional and tumor-modifying programs. In this view, stromal heterogeneity includes a mechanically tunable CAF state driven by volumetric adaptation, reframing fibroblast plasticity as a dynamic, mechanoresponsive feature of the TME. Although direct effects of matrix stiffness on gene regulation are not excluded (e.g., integrin–FAK/Src signaling and downstream RhoA–ROCK mechanotransduction), the prominence of high PDPN-expressing elongated cells across soft and stiff conditions suggests that CAF activation tracks more closely with single-cell morphological state (volume/elongation under 3D confinement) than with bulk modulus alone. Consistent with prior 3D hydrogel study showing that volumetric expansion of CAFs is governed not only by stiffness but also by matrix deformability (e.g., stress relaxation)[19], these results support a model in which ECM mechanics bias CAF state distributions rather than deterministically specifying a single defined phenotype.

Matrix stiffness was associated with a shift in YAP subcellular localization in 3D cultured CAFs. However, single-cell analysis indicated that YAP localization is more closely coupled to the resulting cellular volumetric state than to bulk modulus per se. Immunofluorescence analysis showed that CAFs embedded in soft matrices exhibited higher nuclear-to-cytoplasmic (N/C) YAP ratios than those in stiff matrices (Fig. 3A, B). Single-cell quantification confirmed lower YAP nuclear-to-cytoplasmic (N/C) ratios in soft matrices compared with stiff matrices, despite comparable cell density and staining conditions (Fig. 3C). Across cells, YAP N/C ratios showed substantial heterogeneity within each condition and no strong monotonic dependence on either cell volume or nuclear volume (Fig. 3D,E). In soft matrices, the highest YAP N/C outliers were more frequently observed among smaller-to-intermediate cell volumes, whereas cells at the largest volumes rarely exhibited very high YAP N/C ratios (Fig. 3D). In stiff matrices, YAP N/C ratios similarly spanned a broad range at small-to-intermediate volumes, with considerable overlap with the soft condition and no discernible correlation between cell volume and YAP localization was identified Fig. 3D,E). Associations with nuclear sphericity were weak and not clearly directional (Fig. 3F). Notably, both soft and stiff matrices contained CAF subpopulations with low nuclear YAP, indicating that bulk modulus alone does not fully segregate CAF states and supporting a model in which matrix mechanics and microarchitecture bias single-cell volumetric/morphological states, within which YAP localization varies.

**Fig. 3.**
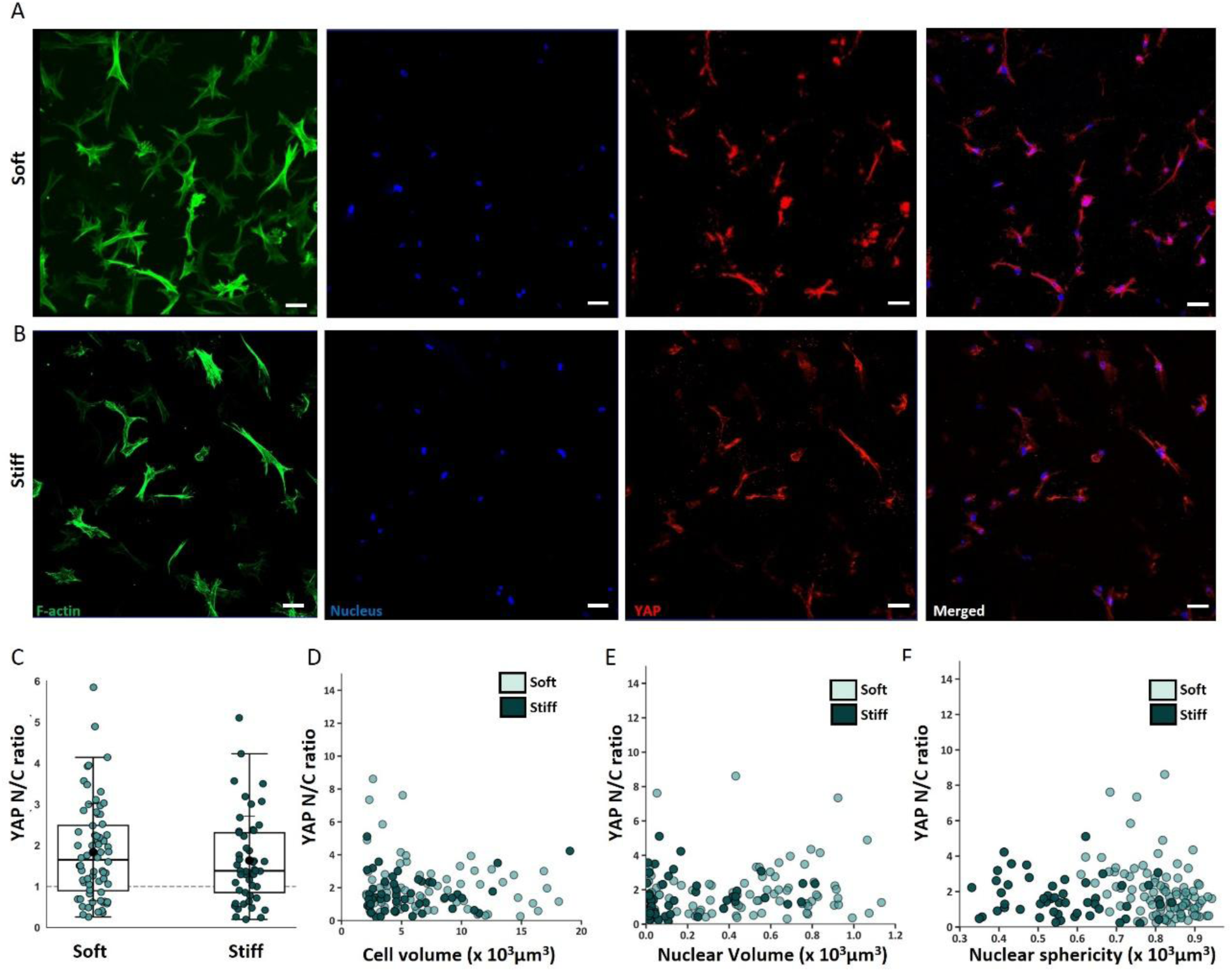
YAP nuclear localization is reduced in stiff 3D matrices and tracks CAF volumetric geometry. A) Representative confocal images of CAFs encapsulated in soft (∼2 kPa) and stiff (∼40 kPa) GelMA matrices and stained for F-actin (green), nuclei (DAPI, blue), and YAP (red). Merged images illustrate condition-dependent intracellular YAP localization. Scale bar, 50 μm. B) Quantification of the YAP nuclear-to-cytoplasmic (N/C) intensity ratio at single-cell resolution shows a reduction trend in stiff versus soft matrices. Box plots indicate median and interquartile range; individual dots represent single cells. C) Scatter plot of YAP N/C ratio versus total cell volume, indicating an inverse relationship across cells from both soft and stiff matrix conditions. (D-F) Scatter plots of YAP N/C ratio versus nuclear geometric parameters (nuclear volume and nuclear sphericity), suggesting that intracellular geometry contributes to YAP localization in 3D.

The observed reduction in nuclear YAP localization in stiff matrices supports a model in which 3D confinement and volumetric restriction suppress YAP nuclear retention in 3D environments [28,29]. In soft matrices, permissive volumetric expansion and more spherical nuclear morphology facilitate YAP nuclear entry, whereas in stiff, constrained cells, nuclear geometry may limit nuclear import or enhance cytoplasmic retention. This behavior contrasts with classical 2D systems, where increased substrate stiffness and cell spreading typically enhance YAP activation, suggesting that volumetric strain and spatial confinement dominate YAP regulation in 3D contexts. This suggests that, unlike 2D systems, where surface spreading drives YAP activation, in 3D contexts, volumetric strain and spatial confinement predominantly regulate YAP mechanotransduction. Overall, these findings contrast with classical 2D systems, where stiffness enhances YAP activation, and instead suggest that in 3D environments, spatial confinement and volumetric geometry play dominant roles in regulating YAP mechanotransduction.

To investigate how paracrine signals from CAFs cultured in soft versus stiff matrices influence cancer cell therapeutic response, we employed a 3D co-culture system in which MCF-7 spheroids were treated with paclitaxel (Fig. S5) in the presence of either soft- or stiff-encapsulated CAF niches using a transwell insert (Fig. 4A). RT–qPCR of MCF-7 spheroids after co-culture showed that those exposed to CAFs encapsulated in stiff matrices upregulated a chemoresistance-associated gene program, including drug efflux *(ABCB1),* survival *(BIRC5),* ECM remodeling *(MMP2),* and growth signaling *(EGFR).* In contrast, spheroids co-cultured with CAFs encapsulated in soft matrices primarily increased expression of the stress-associated cell-cycle regulator *CDKN2A (*p16), without strong induction of efflux or survival genes (Fig. 4C).

**Fig. 4.**
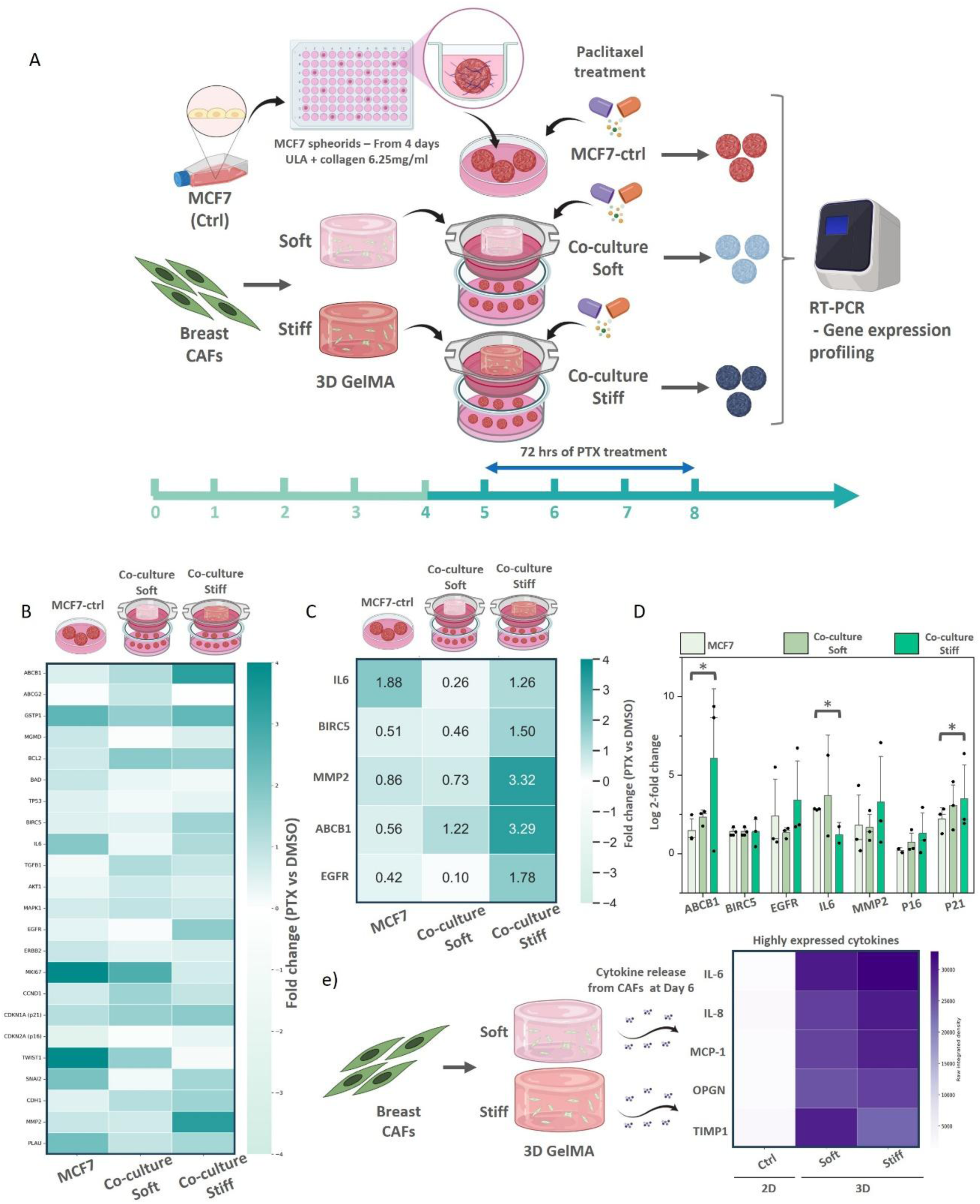
CAFs encapsulated in soft versus stiff matrices differentially induce MCF-7 spheroids toward chemoresistance. A) Schematic of the 3D co-culture and drug-treatment workflow. MCF-7 spheroids were generated in ultra-low attachment (ULA) plates with collagen overlay and co-cultured with breast cancer–associated fibroblasts (CAFs) encapsulated in soft (2 kPa) or stiff (40 kPa) 3D GelMA matrices. Spheroids were treated with paclitaxel (PTX) for 72 h prior to gene expression analysis by RT–qPCR. B) Heatmap showing relative expression of selected drug resistance and survival-associated genes in MCF-7 spheroids cultured alone or co-cultured with CAFs encapsulated in soft or stiff matrices following PTX treatment. Values represent log₂ fold change relative to DMSO-treated controls. C) Quantitative comparison of representative resistance pathways, including drug efflux *(ABCB1),* anti-apoptotic survival *(BIRC5),* extracellular matrix remodeling *(MMP2),* receptor tyrosine kinase signaling *(EGFR),* and stress-associated markers (e.g., p16), across co-culture conditions. D) Bar plots confirming matrix-dependent differences in PTX-responsive gene expression. Data represent mean ± SEM; P < 0.05, P < 0.01 by one-way ANOVA with post hoc testing; n = 3 biological replicates. E) Cytokine profiling of conditioned media collected from CAFs encapsulated in soft and stiff matrices, highlighting enrichment of chemoprotective cytokines (IL-6, MCP-1, OPG, TIMP-1) across conditions.

CAFs are known to influence cancer cells through paracrine mechanisms. In addition to soluble cytokines, CAFs regulate tumor behavior through extracellular matrix remodeling, direct cell–cell interactions, metabolic exchange, and extracellular vesicle transfer. Thus, to understand if stiffness regulates CAF secretion, we measured the cytokine release after culture in soft and stiff matrices. Cytokine profiling of conditioned media showed that CAFs encapsulated in either soft or stiff matrices secreted higher levels of IL-6, MCP-1, OPG, and TIMP-1 than 2D CAF controls (Fig. 4D). Extending this observation, CAFs cultured in 3D matrices exhibited similarly elevated secretion across stiff conditions relative to 2D CAFs, suggesting that dimensionality is a dominant driver of CAF secretory output (Fig. S6). Because the cytokine array was performed as a pilot screen with a single biological replicate (n=1), we interpret these data qualitatively as hypothesis-generating and prioritizing conclusions supported by the functional tumor transcriptional readouts. Moreover, it is important to note that this cytokine profiling was performed at the endpoint and reflects bulk secretion levels, which may not capture transient, spatially localized, or early priming-dependent signaling events. Notably, despite comparable levels of these established chemoprotective factors, only CAF encapsulated from stiff matrices consistently elicited a multi-axis resistance phenotype in tumor spheroids. This discrepancy indicates that chemoresistance is not dictated by bulk cytokine abundance alone but instead reflects a context-dependent CAF niche effect shaped by prior mechanical priming. These findings raise the possibility that stiffness-dependent CAF regulation of tumor cells involves mechanisms beyond soluble cytokine abundance, potentially including matrix remodeling–mediated mechanical feedback, altered cell–matrix architecture, or contact-dependent signaling pathways that are amplified in 3D.

To compare how CAFs encapsulated in 3D soft (vs stiff) GelMA matrices influence drug response in 2D MCF-7 monolayers, we also cocultured the encapsulated CAFs with MCF-7 monolayers. This 2D setup served as a reference to test whether the stromal-driven drug-response signatures observed in 3D spheroids are uniquely enabled by 3D architecture. The transcriptional response to CAF co-culture differed markedly between 2D monolayer and 3D spheroid contexts, with spheroids exhibiting a broader resistance-associated gene program (Fig. 4; Fig. 5).

**Fig. 5.**
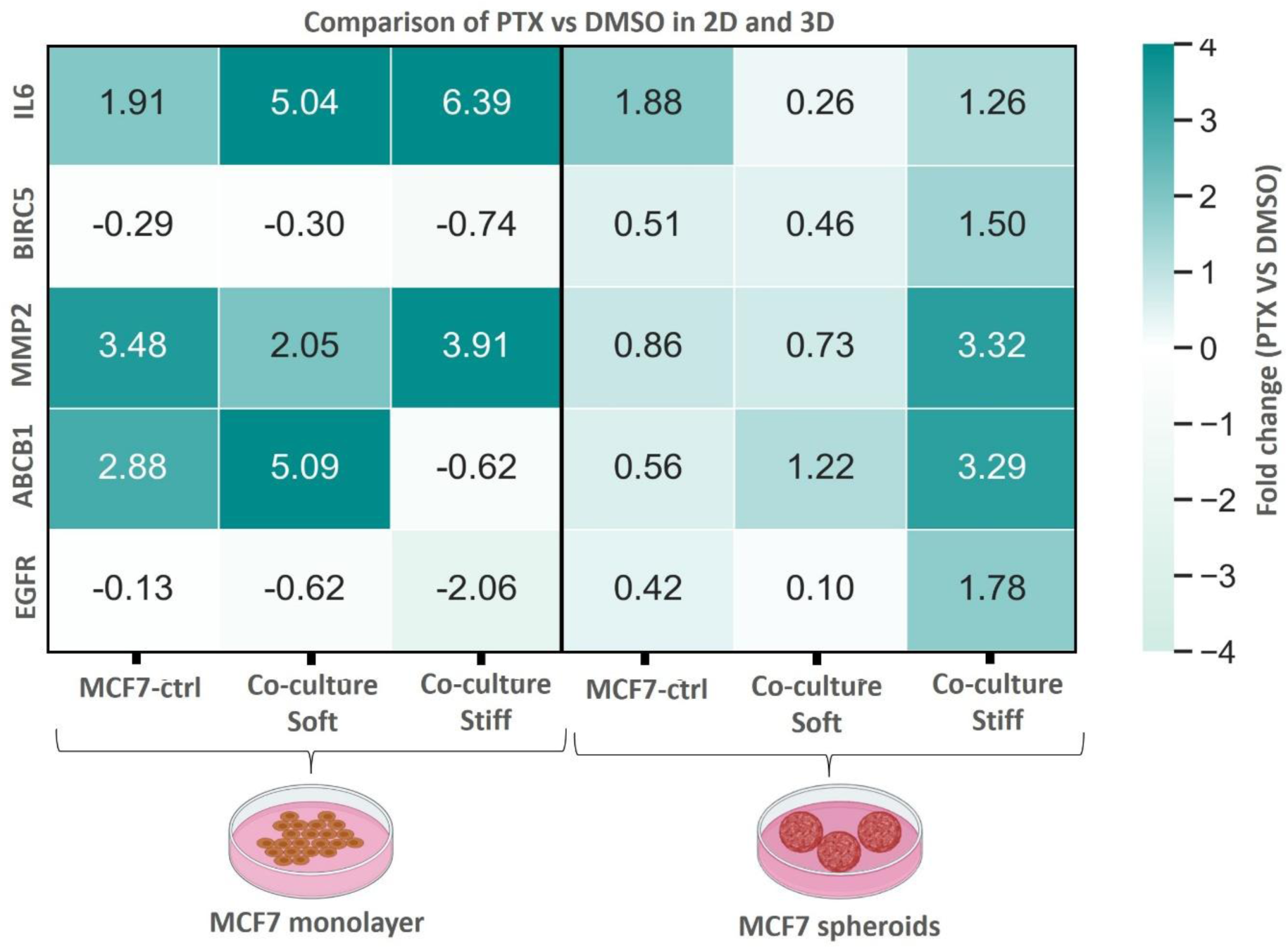
Differential drug-resistance gene responses in MCF-7 monolayers versus spheroids under CAF co-culture. Heatmap showing log₂ fold-changes (PTX vs DMSO) in selected drug-resistance associated genes *(IL6, BIRC5, MMP2, ABCB1, EGFR)* measured by RT–qPCR in MCF-7 cells cultured as 2D monolayers or 3D spheroids. MCF-7 cells were analysed under three conditions: monoculture (Ctrl), co-culture with CAFs encapsulated in soft GelMA matrices (∼2 kPa), or co-culture with CAFs encapsulated in stiff GelMA matrices (∼40 kPa). Gene expression values are normalized to housekeeping genes and reported relative to DMSO-treated controls within each culture format. n = 3–4 independent biological replicates per condition. One-way ANOVA with multiple-comparison correction was applied to ΔCt values. PTX, paclitaxel; CAFs, cancer-associated fibroblasts; DMSO, dimethyl sulfoxide.

In 2D monolayers, CAF co-culture elicits a transcriptional response in MCF-7 cells dominated by IL-6, consistent with an acute stress-associated program. Within this context, CAFs from soft matrices preferentially increase *ABCB1,* suggestive of efflux priming, whereas CAFs induce the strongest IL-6 response but fail to robustly activate efflux (*ABCB1)* or survival-associated genes *(EGFR, BIRC5)* (Fig. S6). These patterns indicate that, in 2D, stromal influence is largely constrained to stress signaling and does not translate into coordinated resistance circuitry. In contrast, when tumor cells are organized as 3D spheroids, CAFs in stiff matrices emerge as a dominant chemoprotective niche, driving concerted upregulation of *ABCB1, MMP2, BIRC5*, and EGFR, accompanied by reduced proliferative signaling *(MKI67)* and only modest IL-6 induction (Fig. 4B–D; Fig. S6).

The selective emergence of this coordinated resistance program in 3D suggests that architectural context enables integration of mechanically conditioned CAF states into stable transcriptional rewiring of tumor cells. The data therefore support a causal sequence in which mechanical priming tunes CAF state, but 3D context is necessary for that state to be functionally integrated into a stable, resistance-promoting tumor niche. Together with earlier volumetric and mechanophenotypic analyses, these results demonstrate that ECM stiffness imprints CAFs with distinct functional programs, and that CAF volumetric state acts as an intermediate linking matrix mechanics to tumor drug-response signatures.

## 4. Discussion

While studies have shown that increased elastic modulus can promote CAF activation [12,32–34], a simple binary classification of “soft” versus “stiff” environments is insufficient to explain why activated CAF programs emerge across mechanically diverse niches, or why CAFs in stiff GelMA matrices can exhibit reduced nuclear YAP a canonical mechanotransduction readout, yet remain transcriptionally and functionally active.

This present study addresses this gap by identifying the CAF volumetric state defined by coupled changes in cell volume, surface area, and shape as a mechanistic intermediary that translates bulk mechanical cues into heterogeneous intracellular signaling and downstream functional responses [17,26,27,35–38]

Importantly, these states are not uniquely determined by bulk modulus. Instead, bulk modulus biases the probability distribution of single-cell volumetric states without imposing a one-to-one mapping between stiffness and state [39]. This framework helps explain why activated CAF programs are observed beyond a strictly “stiff-only” context and highlights CAF heterogeneity as an emergent outcome of mechanically biased yet non-exclusive cell-state occupancy [40–42]

A central implication of our findings is that CAF functional activation aligns more closely with single-cell volumetric geometry than with bulk modulus alone [37,43,44]. PDPN expression, for example, tracked most strongly with CAFs of large volumetric phenotype, regardless of whether those cells resided in soft matrices or stiff matrices (Fig. 2) [45,46]. Conceptually, “volumetric state” is attractive because it operationalizes the integrated cellular consequence of multiple ECM features, elasticity (bulk modulus), pore-scale confinement, and stress transmission into a measurable phenotype that can be mapped at single-cell resolution [35,38,39,47,48]. Consistent with this view, the persistence of multiple morphotypes within each condition suggests that ECM mechanics shape CAF populations by shifting state occupancy rather than enforcing a uniform phenotype [19,20,49].

These geometry-linked states provide a mechanistic lens for interpreting 3D mechanotransduction behavior that diverges from classical 2D paradigms. In 2D, increasing substrate stiffness and cell spreading generally promote YAP/TAZ nuclear localization and transcriptional activation [25,50,51]. By contrast, CAFs encapsulated within stiff matrices exhibited reduced nuclear YAP and increased cytoplasmic retention (Fig. 3).

Notably, YAP nuclear-to-cytoplasmic ratios correlated with cell and nuclear morphometrics, supporting the idea that volumetric confinement and solid stress can override stiffness-dependent YAP activation in 3D contexts [27,29,52]. The presence of YAP-low CAFs in both soft and stiff GelMA matrices further underscores that YAP activity is governed by a cell-scale mechanical state rather than a modulus-gated binary switch [32,53–55]. Despite reduced YAP nuclear localization, CAFs in stiff matrices remained transcriptionally active and matrix-remodeling competent, indicating engagement of a YAP-independent mechanotransduction route [56]. Elevated *MRTF-A* and *TRPV4* expression in CAFs of stiff matrices is consistent with compensation through pathways sensitive to cytoskeletal tension, membrane strain, or volumetric stress (Fig. 3; Fig. S4) [38,57–60]

While our data establish association rather than direct causality, they motivate a working model in which persistent 3D confinement biases mechanotransduction away from canonical YAP-dependent signaling and toward alternative mechanosensing pathways. This model offers a plausible explanation for how CAFs can remain transcriptionally and functionally active in desmoplastic-like, high modulus environments despite suppressed nuclear YAP. More broadly, recent studies on bioengineered hydrogels indicate that elastic modulus, viscoelasticity, and integrin engagement can bias CAF programs at the bulk scale [2,19,61]. Our results extend this view by highlighting volumetric mechanophenotypic states as a single-cell organizing axis that links extracellular mechanics to intracellular signaling and functional output without presupposing predefined molecular subtypes. The functional relevance of this mechanophenotypic heterogeneity was most clearly revealed in tumor co-culture assays. CAFs encapsulated from stiff GelMA matrices induced a multidrug chemoresistance-associated program in MCF-7 spheroids, including coordinated upregulation of drug efflux, survival, growth signaling, and matrix remodeling pathways (Fig. 4)[21–24,62]. In contrast, CAFs from soft GelMA matrices elicited stress-adaptive responses marked by checkpoint activation without robust induction of survival or efflux circuitry [63–65].

Notably, this divergence could not be explained by bulk cytokine abundance alone, as both soft and stiff GelMA matrices induce CAFs to secrete comparable levels of several established chemoprotective factors [66–68] Because our cytokine profiling was performed with limited biological replication (n=1) at endpoint, we treat it as hypothesis-generating and focus our interpretation on concordant patterns rather than stiffness-specific claims.

One plausible explanation is that mechanical conditioning alters the composition, timing, and spatial presentation of stromal signals in 3D, such that tumor cells respond to an integrated, mechanically imprinted niche rather than responding to any single cytokine in isolation [20,49]. In addition, CAFs from stiff GelMA matrices, may regulate tumor transcription through mechanisms not captured by endpoint conditioned-media assays, including ECM remodeling mediated mechanochemical feedback, extracellular vesicle cargo, and contact- or adhesion-dependent signaling that is amplified by 3D architecture [33,69–74] Together, these results are consistent with a sequence in which mechanical conditioning shapes CAF state, while 3D tumor organization facilitates its integration into a stable, resistance-promoting niche [21,75,76]

Several limitations should be considered. All experiments were conducted using primary breast CAFs in GelMA-based 3D matrices, and it remains to be established whether identical volumetric rules generalize across CAF subtypes, tumor origins, or ECM compositions [8,18,41,42,77–79]. In our system, bulk modulus and microarchitecture were co-varied, limiting the ability to fully decouple elastic modulus from pore-scale confinement and stress transmission[80–82]. Mechanotransducive activity was inferred from localization and expression markers rather than direct force readouts or ion-flux measurements [38,83,84]. A key limitation is that the cytokine profiling was based on a pilot array (n=1) at an endpoint timepoint; therefore, follow-up validation using targeted ELISA/Luminex (e.g., IL-6, MCP-1, OPG, TIMP-1) with ≥3 independent CAF donors/replicates and time-resolved sampling will be required to quantify stiffness-dependent differences and capture transient priming dynamics. Finally, the temporal stability, reversibility, and potential hysteresis (“memory”) of volumetric states were not directly tested, and time-resolved profiling will be needed to identify early priming-dependent signaling events [85–87] Definitive causal inference will require perturbations that selectively modulate cell volume or confinement under constant [87–89].

Overall, this study reframes stromal targeting strategies by shifting emphasis from macroscopic stiffness modulation toward control of volume- and geometry-regulating mechanoadaptation. Tumor softening alone may be insufficient if CAFs remain volumetrically confined and mechanically activated through alternative mechanotransduction pathways [90,91]. Instead, targeting mechanosensing nodes that regulate volumetric adaptation, such as *TRPV4* and the actin *MRTF* axis may provide a more precise approach to stromal normalization[58,92–94].

We propose the following experimentally tractable next steps to test causality and therapeutic leverage: (i) perturb *TRPV4* in CAFs (pharmacologic inhibition and/or siRNA/CRISPR knockdown) and determine whether CAF volumetric-state distributions and downstream spheroid chemoresistance signatures (ABCB1, BIRC5, EGFR, MMP2) are reduced during paclitaxel treatment; (ii) orthogonally perturb *MRTF/SRF* signaling (e.g., *CCG-203971* or *MRTFA* knockdown) to assess whether YAP-independent activation and chemoprotective niche behavior require the actin–*MRTF* transcriptional axis; and (iii) decouple stiffness from confinement by engineering matrices with matched elastic modulus but altered mesh size and/or viscoelasticity (e.g., varying GelMA formulation or incorporating stress-relaxing elements) to test whether volumetric restriction, rather than stiffness per se, better predicts pathway engagement and tumor protection [82,92,95]. In parallel, a time-resolved secretome profiling (early vs late time points) would directly test whether mechanical context drives transient cytokine dynamics that are not captured by endpoint conditioned-media assays, and whether these dynamics better align with resistance acquisition.

In conclusion, our findings establish CAF volumetric state as an organizing principle linking ECM mechanics, mechanotransduction pathway selection, and tumor drug response, providing a mechanistic foundation for precision targeting of the TME without indiscriminate stromal depletion.

## CRediT authorship contribution statement

**Somayadineshraj Devarasou:** Conceptualization, Methodology, Data curation, Formal analysis, Investigation, Writing – original draft.

**Nam Ji Sung:** Methodology, Formal analysis, Data curation.

**Seok Hoon Ham:** Methodology, Software, Data curation.

**Martin Kiwanuka:** Resources, Methodology, Writing – review & editing.

**Jennifer L. Young:** Resources, Investigation, Writing – review & editing.

**Jennifer H. Shin:** Supervision, Conceptualization. Investigation, Funding acquisition, Writing – review & editing.

## Acknowledgements

This research was supported by a grant of the Korea-US Collaborative Research Fund (KUCRF), funded by the Ministry of Science and ICT and Ministry of Health & Welfare, Republic of Korea (to J.H.S by grant number: RS-2024-00468873) and the Ministry of Education under the Research Centres of Excellence programme through the Mechanobiology Institute at the National University of Singapore and the Biomedical Engineering Department at the National University of Singapore (to J.L.Y.).

The authors thank Dr. Yongsung Hwang (Soonchunhyang University) and Dr. Khoon Lim (University of Sydney) for guidance regarding the GelMA and Ru/SPS hydrogel synthesis. The authors also thank the Fluid and Interface Laboratory, Department of Mechanical Engineering, KAIST, for assistance with hydrogel rheological characterization, and the Neutron Scattering Laboratory, Department of Nuclear and Quantum Engineering, KAIST, for support with the GelMA freeze-drying process.

## Supplementary Figures

**Fig. S1.**
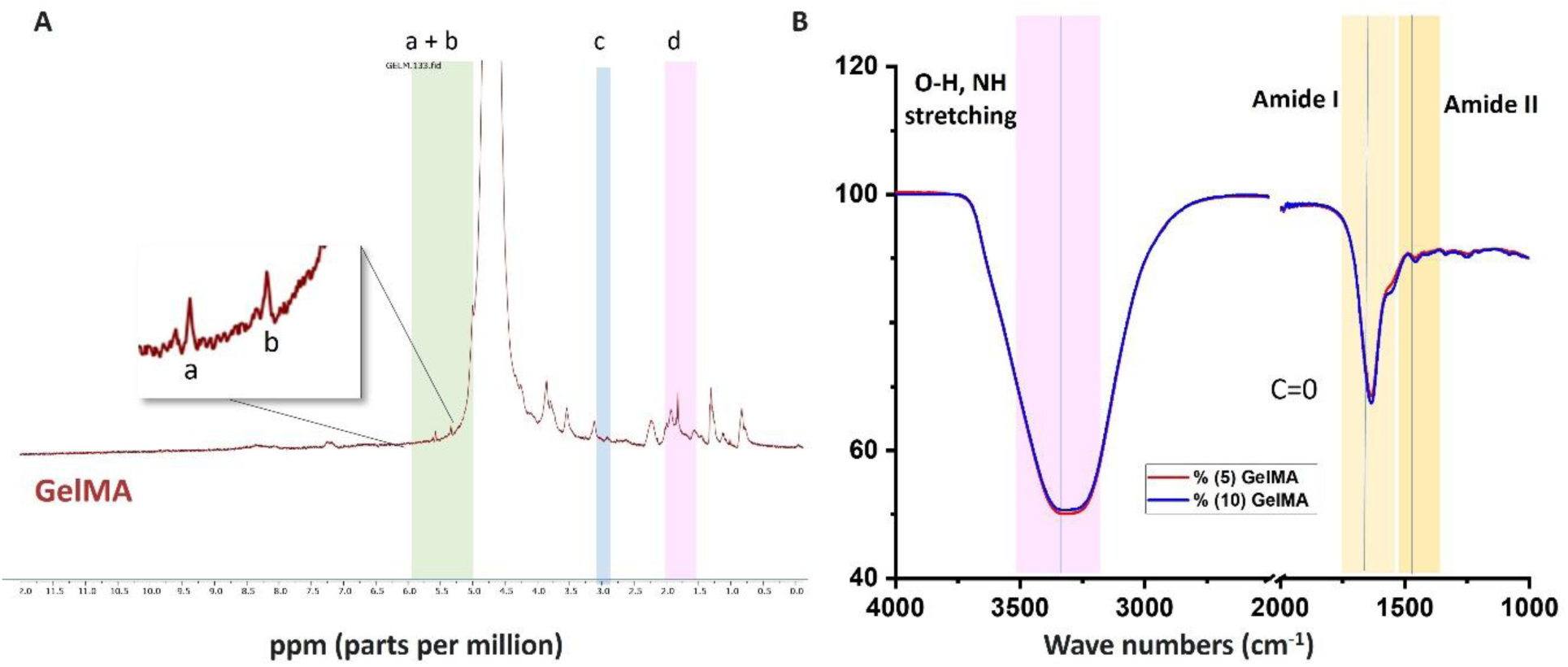
Spectroscopic validation of gelatin methacryloyl (GelMA) synthesis. A) Representative ^1H NMR spectrum of GelMA. Peaks corresponding to acrylic protons of the methacrylate group (a + b) are observed in the 5–6 ppm range. Signals at ∼2.9 ppm correspond to methylene protons of lysine residues in the gelatin backbone (c), while methyl protons of the methacryloyl groups are detected at ∼1.5–2.0 ppm (d), indicating successful incorporation of methacrylate moieties. B) FTIR spectra of gelatin and GelMA. GelMA exhibits characteristic amide I and amide II bands, along with a C=C stretching vibration near ∼1635 cm⁻¹, consistent with methacrylation of gelatin. A broad absorption band in the 3200–3400 cm⁻¹ range corresponds to O–H stretching associated with hydrogen bonding and residual water within the hydrogel. Representative spectra from ≥3 independent synthesis batches; GelMA, gelatin methacryloyl; FTIR, Fourier transform infrared spectroscopy.

**Fig. S2.**
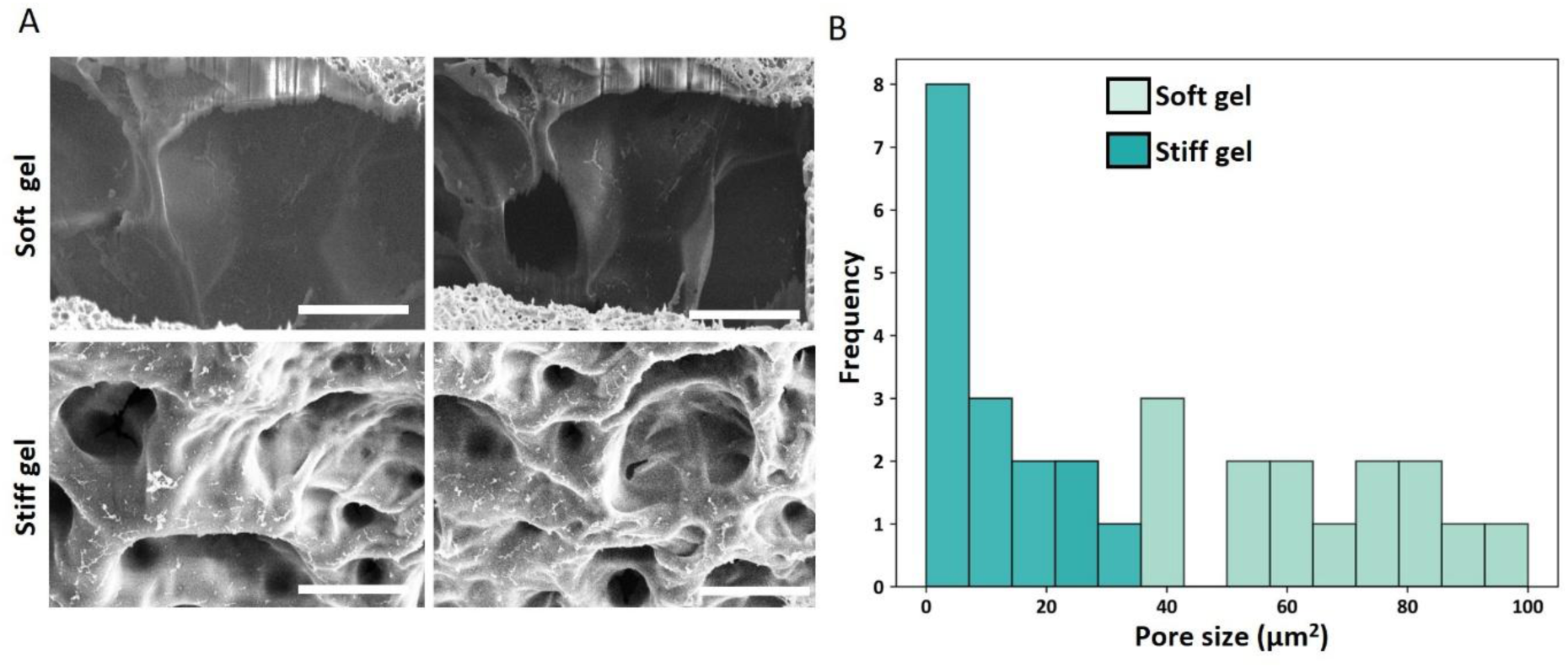
Microstructural characterization of GelMA hydrogels across soft and stiff matrix conditions. A) Representative scanning electron microscopy (SEM) images showing the internal pore architecture of soft (∼2 kPa) and stiff (∼40 kPa) GelMA hydrogels. Soft gels exhibit larger and more heterogeneous pores, whereas stiff gels display a denser network with reduced pore size and increased structural compactness. Scale bar, 5 μm. B) Quantitative pore size distribution analysis of soft (2 kPa) and stiff (40 kPa) GelMA hydrogels. Histograms show a right-shifted distribution in soft gels, indicating larger pore sizes, whereas stiff gels predominantly exhibit smaller pore dimensions.

**Fig. S3.**
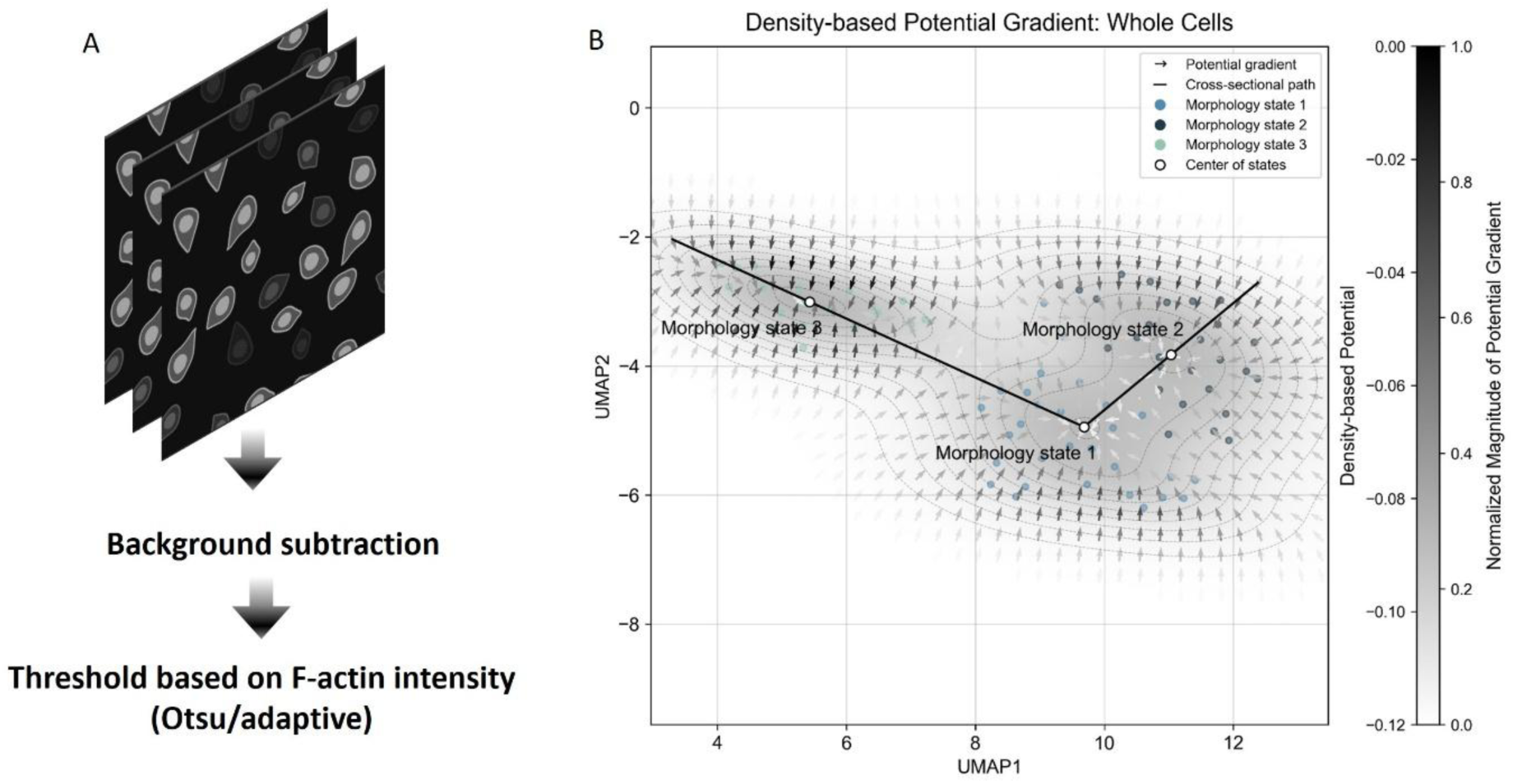
Image-processing and density-based morphometric mapping of CAF volumetric states. A) Schematic of the image-processing workflow applied to confocal z-stacks of CAFs stained for F-actin. Raw images were subjected to background subtraction followed by adaptive thresholding based on F-actin intensity (Otsu method) to generate segmented single-cell masks. Representative segmented projections are shown for CAFs cultured soft and stiff matrices. B) Density-based potential gradient map computed from single-cell morphometric features embedded in UMAP space. Vector fields represent local gradients of the cell-density distribution across the morphospace, revealing continuous transitions between morphological states.

**Fig. S4.**
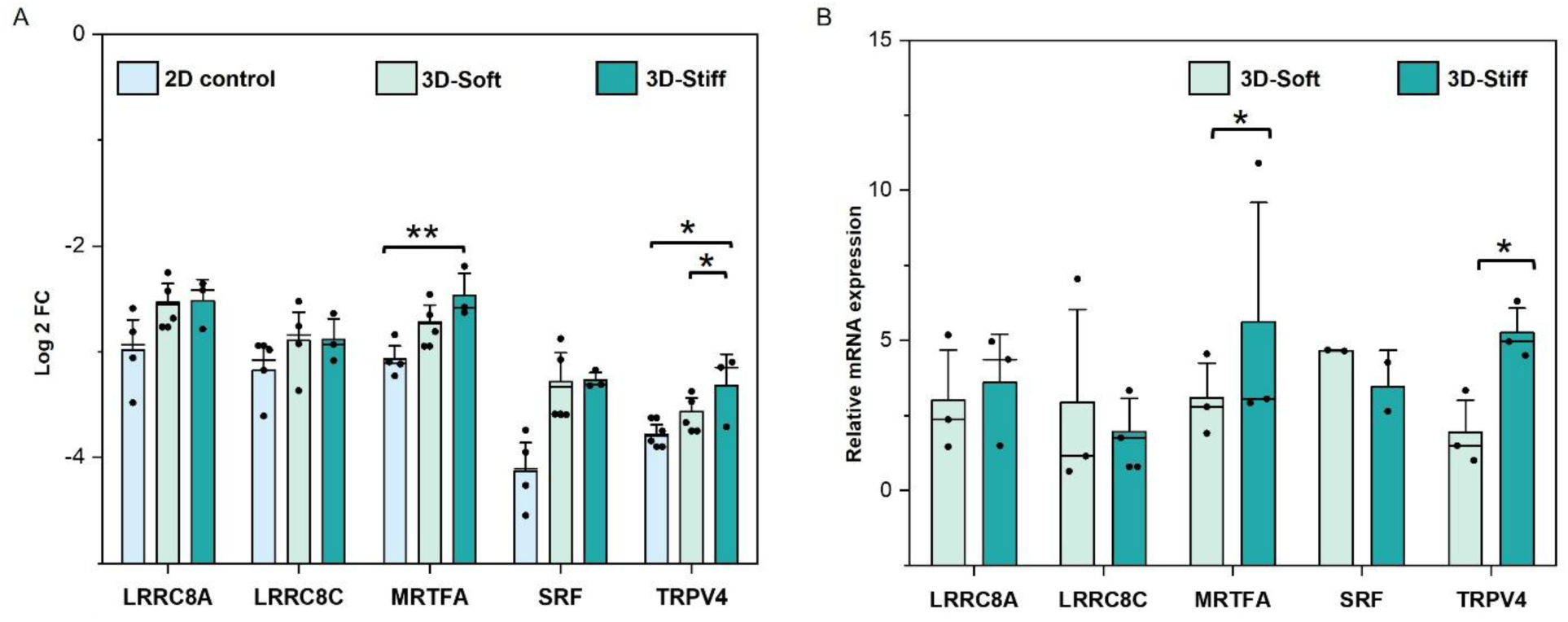
Differential expression of mechanosensing-associated genes in CAFs cultured in soft versus stiff 3D matrices. A) Represent the Log₂ fold-change in mRNA expression of LRRC8A and LRRC8C (volume-regulated anion channels), MRTFA and SRF (actin–transcriptional regulators), and TRPV4 (mechanosensitive ion channel) in CAFs cultured in 3D soft (∼2 kPa) and 3D stiff (∼40 kPa) GelMA matrices, normalized to 2D CAF controls. Individual data points represent biological replicates. B) Relative mRNA expression of the same gene set comparing CAFs cultured in 3D soft (∼2 kPa) versus 3D stiff (∼40 kPa) GelMA matrices, normalized to GAPDH. CAFs in stiff matrices show significantly increased expression of MRTFA and TRPV4 relative to CAFs in soft matrices, whereas LRRC8A, LRRC8C, and SRF show no significant differences. Gene expression was quantified by RT–qPCR and analyzed using the 2^−ΔΔCt^ method. n = 5 independent biological replicates per condition. One-way ANOVA with multiple-comparison correction and unpaired two-tailed Student’s t-test (*p < 0.05, **p < 0.01). Data represent mean ± SEM

**Fig. S5.**
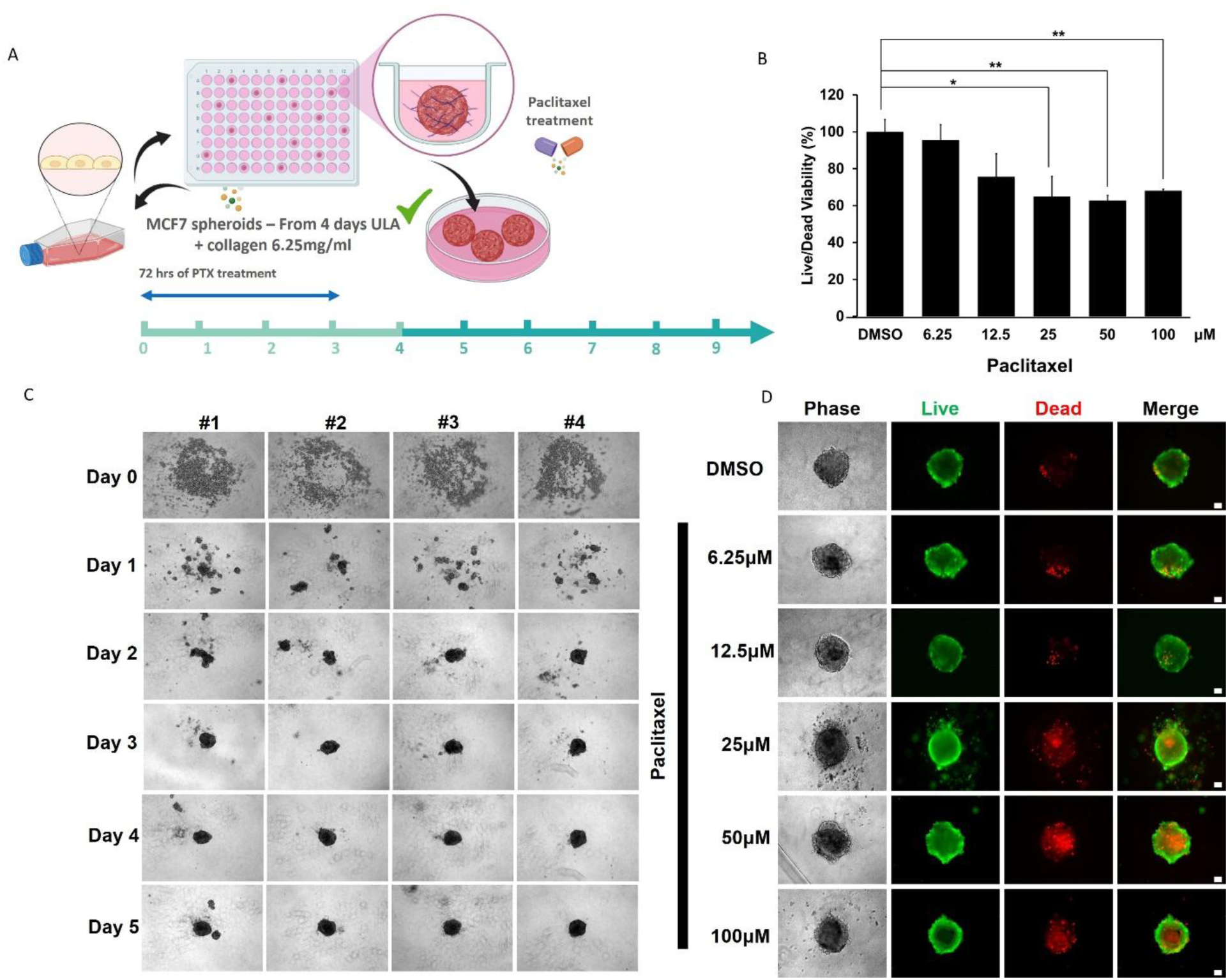
Establishment of MCF-7 spheroid growth and paclitaxel dose selection in ultra-low attachment culture. A) Schematic illustrating the experimental workflow for MCF-7 spheroid formation in ULA plates supplemented with collagen, followed by paclitaxel treatment for 72 h. B) Quantification of MCF-7 spheroid viability following treatment with increasing concentrations of paclitaxel (6.25–100 μM), normalized to DMSO-treated controls. Data demonstrate a dose-dependent reduction in viability, with partial but incomplete loss of viability at 6.25 μM. C) Representative brightfield images showing temporal progression of MCF-7 spheroid morphology over 5 days in ULA culture across independent wells (#1–#4), demonstrating consistent spheroid formation and compaction. D) Representative live/dead fluorescence images of MCF-7 spheroids after 72 h of paclitaxel treatment at the indicated concentrations. Live cells are shown in green and dead cells in red. n = 8 independent spheroids per condition: representative images shown. Mean ± SD; PTX, paclitaxel; ULA, ultra-low attachment; DMSO, dimethyl sulfoxide.

**Fig. S6.**
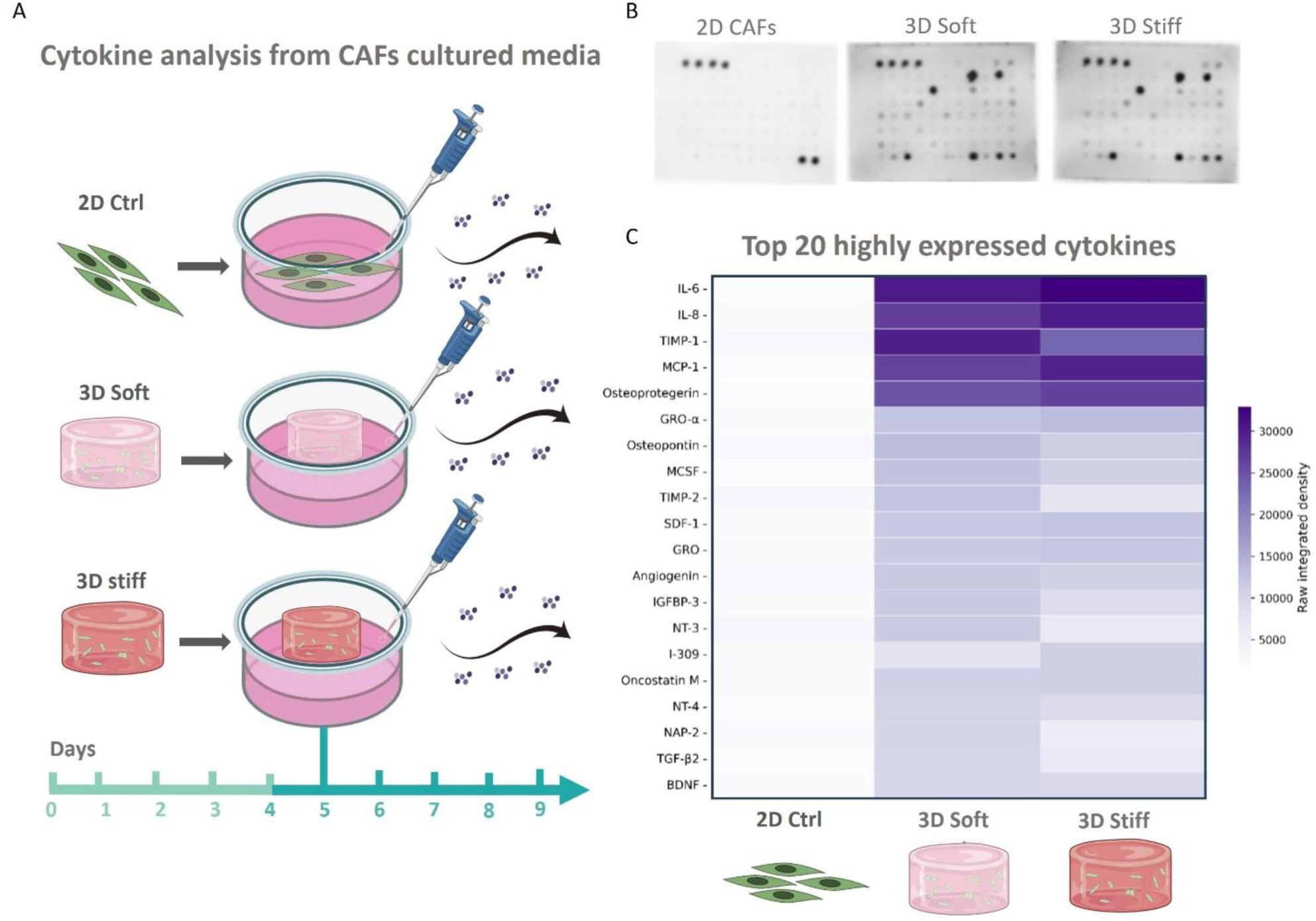
Cytokine profiling of conditioned media from 2D CAFs and CAFs encapsulated in soft versus stiff GelMA matrices. A) Schematic illustrating the experimental workflow for cytokine analysis. Conditioned media were collected from 2D CAF cultures and from CAFs encapsulated in 3D soft (∼2 kPa) and 3D stiff (∼40 kPa) GelMA matrices over the indicated culture period prior to analysis. B) Representative cytokine antibody array membranes showing relative signal intensities for secreted factors in conditioned media from 2D CAFs and CAFs encapsulated in 3D soft and stiff matrices. C) Heatmap summarizing the top 20 highly expressed cytokines across conditions, quantified from antibody array signal intensities. Cytokine levels are displayed as mean normalized intensity values for each condition. Elevated secretion of IL-6, MCP-1, OPG, and TIMP-1 is observed in both 3D soft (∼2 kPa) and 3D stiff (∼40 kPa) CAF cultures relative to 2D CAF controls. n = 1 independent biological replicate per condition (pilot screen).

**Fig. S7.**
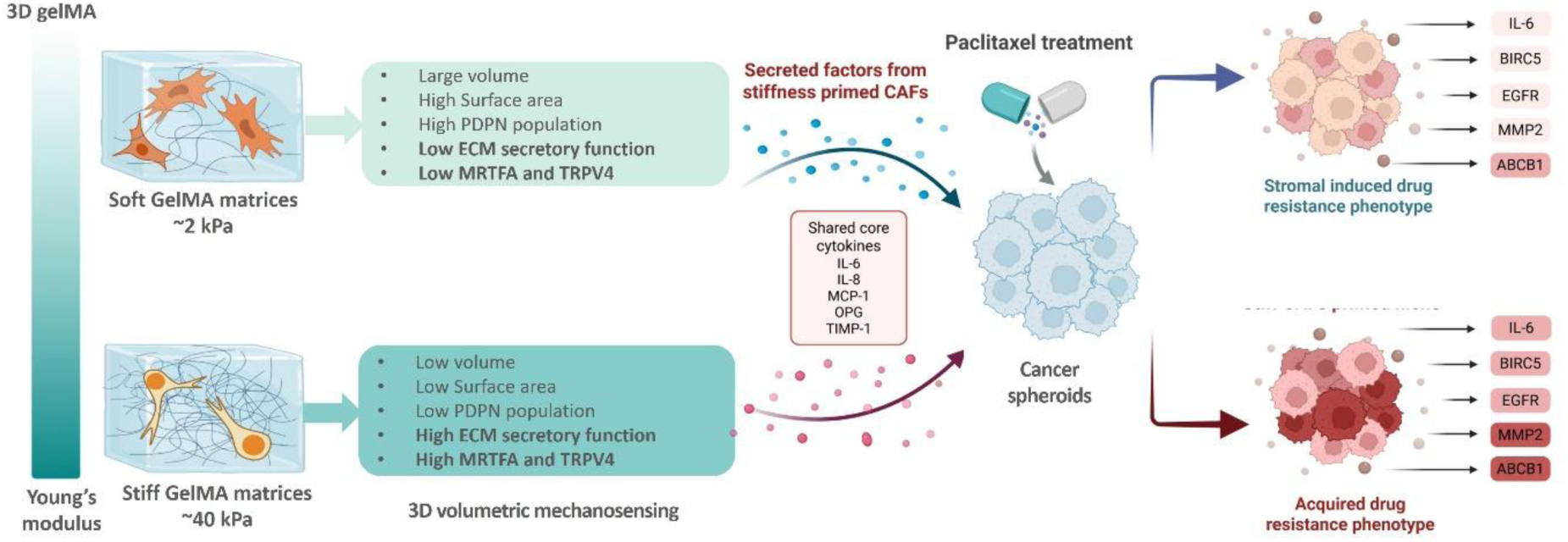
Schematic summary of the working model derived from this study. ECM elastic modulus in 3D GelMA soft versus stiff matrices biases CAF volumetric states, generating distinct mechanophenotypes characterized by differences in cell volume, surface area, morphology, and PDPN expression. Soft (∼2 kPa) matrices favor larger-volume CAFs with expanded surface area, whereas stiff (∼40 kPa) matrices promote more confined, compact, lower-volume morphologies. Both CAF states secrete a shared core cytokine set (IL-6, IL-8, MCP-1, OPG, TIMP-1); however, CAFs encapsulated in stiff matrices create a niche that drives a coordinated, multi-gene chemoresistance program in MCF-7 spheroids following paclitaxel treatment, marked by elevated ABCB1, MMP2, BIRC5, and EGFR expression. In contrast, co-culture with CAFs encapsulated in soft matrices elicits a more limited stromal-induced stress response. The schematic illustrates the proposed volumetric mechanotransduction framework linking 3D matrix mechanics (bulk stiffness and associated confinement), CAF state, and tumor drug response.

**Fig. S8.**
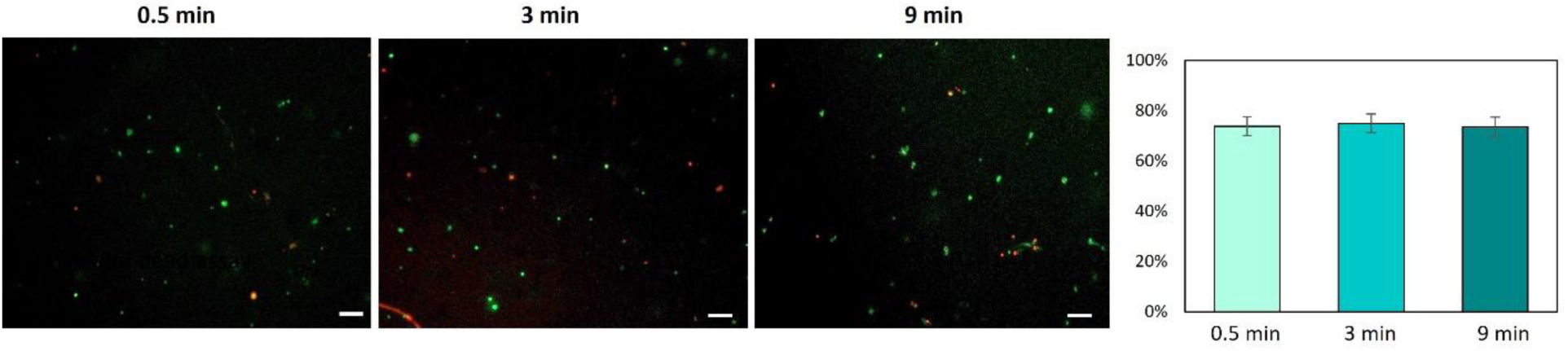
Live/dead staining confirms preserved viability across photo-crosslinking exposure durations. Representative fluorescence images of cells following 0.5, 3, and 9 min light exposure (Ru/SPS crosslinking conditions) show live cells in green and dead cells in red at each time point. Quantification (right) indicates comparable viability across all exposure durations, with no significant decrease observed up to 9 min. Scale bars, 100 µm.

